# Identification of genomic regions implicated in susceptibility to *Schistosoma mansoni* infection in a murine genetic model (backcross)

**DOI:** 10.1101/2022.06.06.494967

**Authors:** Juan Hernández-Goenaga, Julio López-Abán, Adrián Blanco-Gómez, Belén Vicente, Francisco-Javier Burguillo, Jesús Pérez-Losada, Antonio Muro

## Abstract

High levels of infection and severe liver fibrosis in schistosomiasis appear only in a few cases of infected people with high susceptibility. Tissue damage is caused by the inflammatory response to eggs trapped in the liver. The genetic background influences susceptibility to schistosome infection. To assess the genetic basis of susceptibility to schistosomiasis and identify the chromosomic regions involved, we used a backcross strategy to generate a mouse cohort with high variation in schistosomiasis susceptibility. Thus, we crossed the resistant C57BL/6 mouse strain with the susceptible CBA one; and later; the F1 females (C57 x CBA) were backcrossed with CBA males generating the F1BX cohort. The spectrum of phenotypes in the F1BX mice showed gradation from mild to severe disease, lacking a fully resistant group. We differentiated four levels of infection intensity using cluster and principal component analyses and K-means based on parasitological, pathological and immunological trait measurements. Mice were massively genotyped with 961 informative SNPs. We identify 19 new quantitative trait loci (QTL) associated with phenotype indicators of parasite burden, liver lesion, white blood cell populations, and antibody responses evaluated in the backcross cohort. Two QTLs on chromosomes 15 and 18 were simultaneously linked to the number of granulomas, grade of liver lesion and IgM levels. The human syntenic regions are located in chromosomes 8 and 18. None of the significant QTL identified coincided with previously reported mice association or were syntenic with human chromosomes.

**Author summary:** High number of cases of infection and high levels of infection and liver fibrosis in schistosomiasis appear only in a few cases of people with high susceptibility, because the genetic background influences this susceptibility. To assess this, we used a backcross strategy crossing a resistant strain of mouse C57BL/6 with another susceptible CBA, and later F1 females were backcrossed with males susceptible, generating the F1BX cohort, which showed gradation from mild to severe disease. We differentiated four levels of infection according to cluster, principal component analysis and K-means, and based on parasitological, pathological and immunological measurements. Mice were massively genotyped and 19 quantitative trait loci (QTL) were identified, associated with phenotype indicators evaluated in backcross cohort. Two QTLs on chromosomes 15 and 18 were simultaneously linked to the number of granulomas, grade of liver lesion and IgM levels. None of the significant QTL identified coincided with previously reported mice association or were syntenic with human chromosomes.

## Introduction

Schistosomiasis is a widespread parasitic disease caused by trematodes of the genus *Schistosoma* in tropical areas. It is estimated that more than 200 million people worldwide are infected with *Schistosoma* spp., which caused more than 1.4 million DALYs (Disability Adjusted Life Years) in 2017 [1]. Schistosomiasis is a complex disease resulting from the parasite’s interaction with endogenous host factors: such as intensity and duration of infection, nutritional status, and other associated diseases [2]. Significant heterogeneity is observed in everyone’s response to the parasite in the endemic areas. Only a minority of patients suffer severe hepatic fibrosis or had high infection burden [3]. According to SNPs and microsatellite marker studies, a chromosomal region identified in endemic areas of Brazil and Senegal identified the SM1 (*S. mansoni* 1) locus linked to high prevalence and high infection burden in particular family groups [4, 5]. The SM1 susceptibility locus is localized in the long arm of human chromosome 5 (5q31-33). Another locus associated with severe schistosomiasis is called SM2 (*S. mansoni* 2), related to the development of liver fibrosis and located on the long arm of human chromosome 6 (6q22-23) [6, 7]. Despite the reports suggesting genomic regions associated with high intensity of the infection and severe liver damage no compressive information about the genomic regions involved in these processes. Knowledge of molecular markers linked to susceptibility could be helpful to develop protocols for diagnosis and treatment against severe schistosomiasis and enable efficient control measures to prevent infection in more susceptible populations.

Identifying genetic markers of complex diseases, such as schistosomiasis is difficult in the human population. It requires population studies with many cases and controls and a great expense of time and resources without guaranteeing success. However, syngeneic mouse strains are genetically and phenotypically homogenous in schistosomiasis susceptibility and other traits [8].Also, crosses between syngeneic mouse strains generate cohorts of individuals with a tremendous variation in infection susceptibility, which allows the identification of genetic regions, such as QTLs (Quantitative Trait Loci) and expression QTLs (eQTLs), associated with any quantitative trait variation; thus, involved in the resistance or susceptibility to schistosomiasis [9]. Identifying these genetic loci linked to the disease susceptibility will help elucidate the molecular basis of this complex disease [10, 11]. Indeed, linkage studies with SNPs (single nucleotide polymorphisms) have identified chromosomal regions associated with the genetic influence of schistosomiasis and its wide variability in severity. It has been possible to delimit genetic regions involved both in the intensity of the schistosome infection and in the degree of liver fibrosis induced in an individual [12–14]. In mice, laboratory infections with *S. mansoni* lead to granulomatous lesions in the intestine and liver. We carried out, a preliminary study with five syngeneic mice strains (C57BL/6J, FVB/NH, DBA/2J, BALB/c and CBA/2J) and we identified the two most divergent strains in terms of susceptibility to infection, which were CBA/2J more susceptible and C57BL/6J less. The susceptibility to infection in the F1B6CBA hybrid mice was between these two parental strains yielding intermediate susceptibility [15]. The strain differences in the response to *S. mansoni* also offer the chance to study the genetic factors influencing whether an animal can develop intense or mild worm recovery or lesions to this infection. The evidence of differences between more and less susceptible strains led us to adopt a strategy of backcrossing of C57BL/6J onto the CBA background. This approach should generate mice with high phenotypic variability between the two strains.

The general objective of this work was to identify chromosomal regions that carry low penetrance or modifiers genes associated with the susceptibility to infection by *S. mansoni*, by a backcross murine genetic model, with the following specific objectives: characterize the susceptibility to infection by *S. mansoni* defining different disease patterns; to study the association between the pathophenotypes and the related parasitological, pathological and immunological traits and to identify the genomic regions (QTL) linked to the intermediate phenotypes defined after infection with *S. mansoni*.

## Results

### There was a different susceptibility to schistosomiasis in mice related to the genetic background but not to mouse sex

We had previously studied the susceptibility/resistance to *Schistosoma mansoni* infection in C57BL/6J, CBA/2 mouse strains and the F1B6CBA [15]. Now we generated a backcross (F1BX) cohort of 105 mice to identify associated chromosomal regions with *S. mansoni* infection. The F1BX cohort presented few male and female counts after perfusion than those of the F1B6CBA and similar numbers to the resistant parental strain. Regarding the number of eggs per gram of liver, the mean was like the F1B6CBA cohort and intermediate to the parental strains. The number of granulomas in the F1BX cohort was higher than that in the F1B6CBA and similar to the susceptible CBA/2J parental strain (Supplementary Figure 1).

When comparing F1BX males and females we did not observe significant differences in number of recovered male worms, eggs trapped in tissues, fecundity, number of granulomas or injured liver surface (Supplementary Figure 2). Moreover, we observed a wide variability between data collected for different mice of the F1BX varying between resistant C57BL/6J and susceptible CBA/2J strains (Supplementary figure 3).

### Parasitological, pathological and white blood cell populations correlated in the F1BX cohort

Most of parasitological and pathological variables showed direct association between them, but the number of recovered males and females, and worm fecundity showed an inverse association. The hepatic damage variables (granulomas and affected liver surface) the number of peripheral blood lymphocytes at week nine post infection, and the number of eggs in tissues showed an inverse correlation with splenocyte subpopulations. The parasitological and pathological variables were directly associated with immunoglobulin serum levels. The white blood cell subpopulations showed a direct association between them, but not with immunoglobulin levels except B220^+^ splenocytes and IgM, which showed an inverse association. Finally, immunoglobulins showed a direct association between them, unless IgG2a with IgG nor IgM (Figure 1).

**Figure 1.**
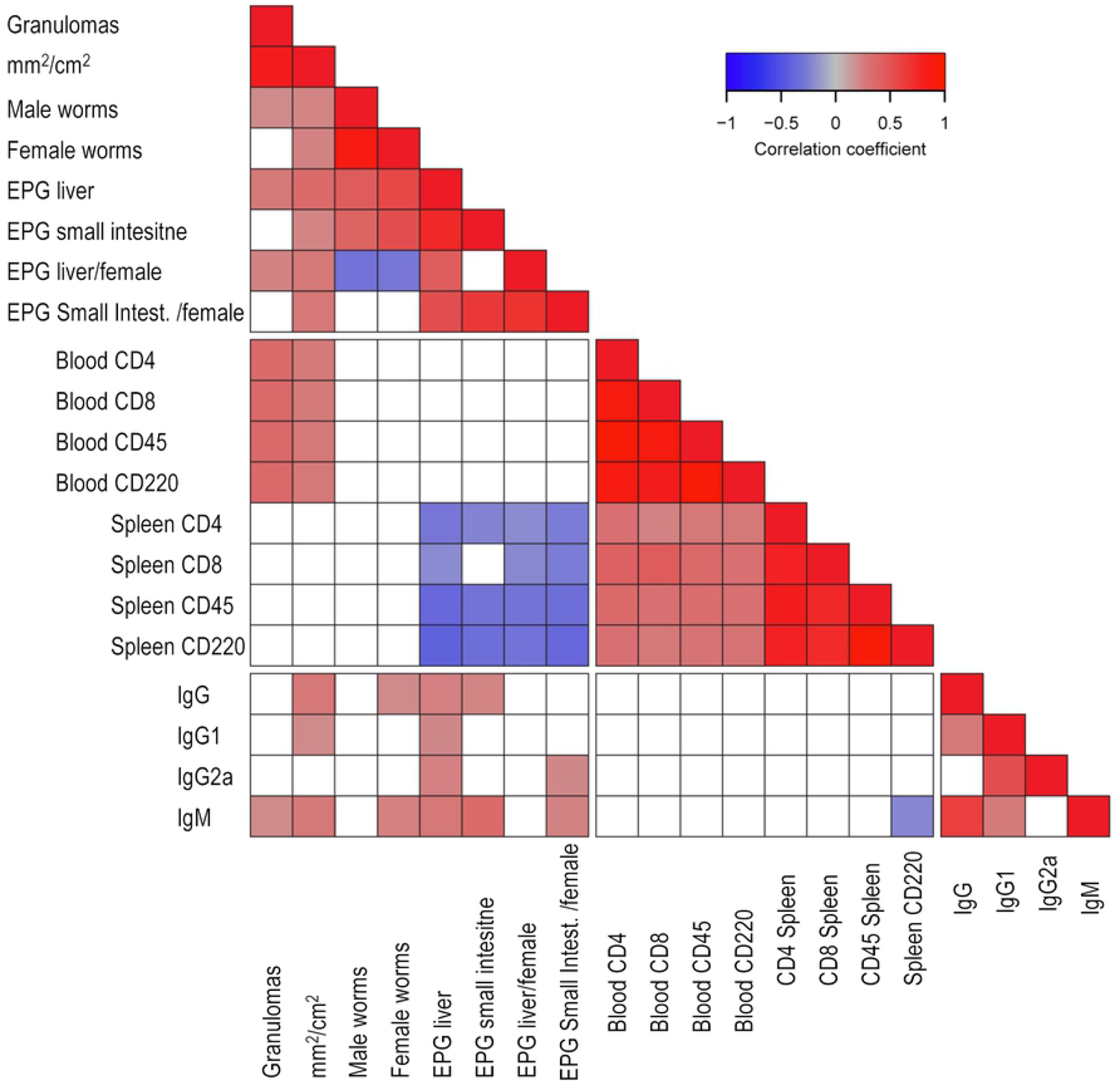
Correlations analysis between variables. Direct correlations are represented in red and indirect correlations are in blue. The intensity of the color represents the correlation coefficient according to the scale. Only those significant correlations were represented (p <0.05). The correlations between variables that were not statistically significant are shown in white (p> 0.05).

### Identification of different degrees of schistosomiasis disease in the F1BX mouse cohort

Multivariant analysis techniques were used to group the mice and analyze the variables’ influence. Initially a clustering analysis (CA) was performed to study similarities in the F1BX cohort using recoveries of male and female worms, eggs in the liver and intestine, fecundity, granulomas, and surface liver damage. Distances between the cases represented in the dendrogram as branches were interpreted as different groups. Data of the F1BX cohort ranging from the more to the less susceptible mouse pointed to four groups, but there were atypical individuals, and it was not easy to establish a threshold. (Supplementary figure 4).

Then, we applied the principal components analysis (PCA) to condense information into two dummy variables and to know the load of each variable. The principal component 1 (PC1) captured 42% of the information and PC2 28%. Then, we shorted our F1BX cohort into four groups in agreement with results obtained using cluster analysis (supplementary figure 5A). Moreover, we observed that all variables influence the PC1 in the same sense. On the other hand, male and female count loads have opposite influences on fecundities in PC2. Eggs in tissues and granulomas had a neutral influence in PC2 (supplementary figure 5B).

Clusters by k-means algorithm need to assign initial mean values for each of the four groups: therefore, we used those obtained from PCA. After the iterations, the algorithm joined the cases around the centroids of each group. In Figure 2, centroids and cases are connected with radios. Mice were grouped in each cluster based on the severity of the infection from mild (Group 1) to severe (Group 4).

**Figure 2.**
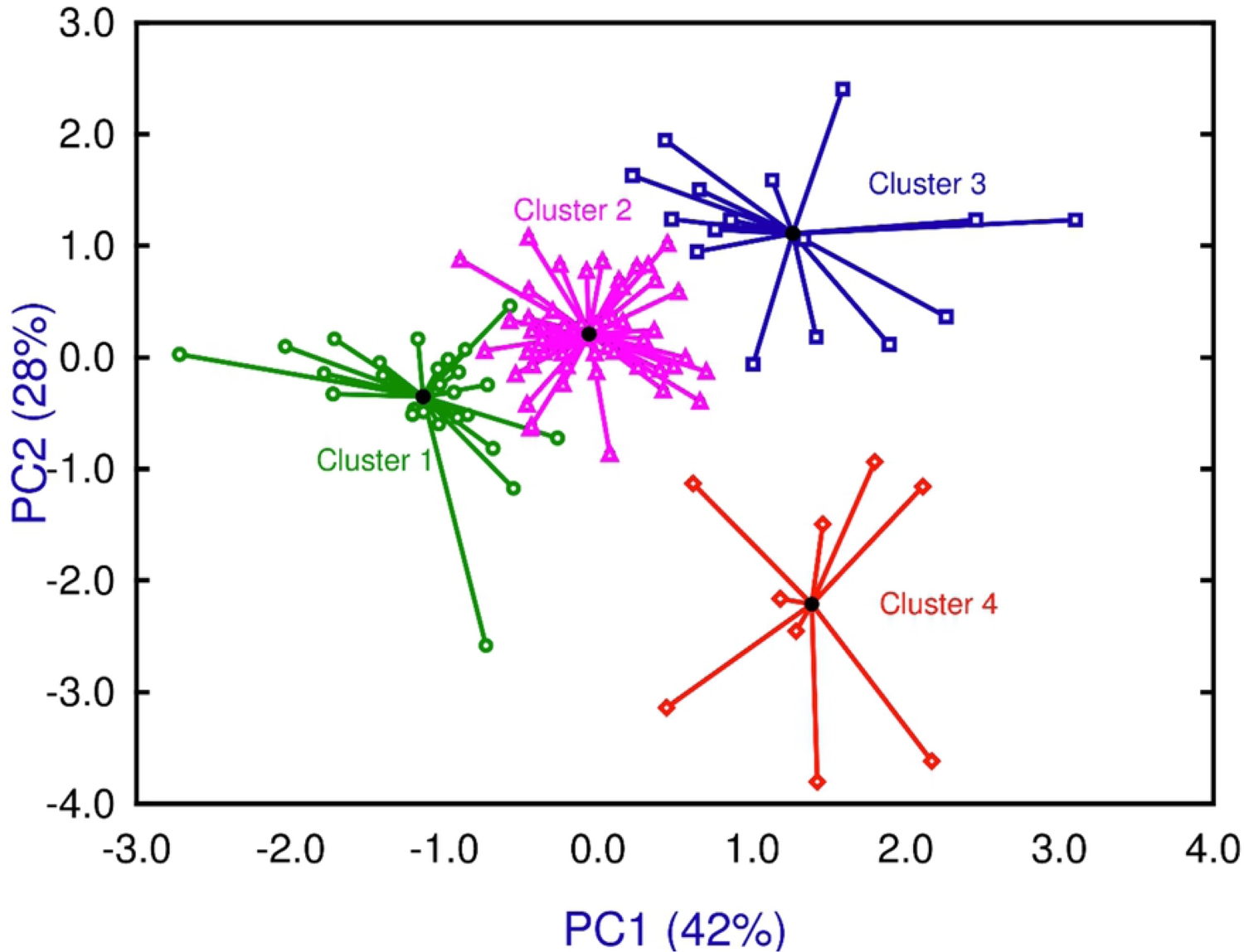
K-means clusters showing four cluster around their centroids ranging severity in four levels from mild (Group 1) to severe (Group 4).

### Pathophenotypic variation in the F1BX cohort in the clusters identified

We evaluated the distribution of each schistosomiasis pathophenotypes and intermediate phenotypes in the four clusters identified in the F1BX cohort by the multivariant analysis. We considered male and female recoveries, eggs in the liver and intestine, granulomas and affected liver area (Figure 3). Also, we studied lymphocyte subpopulations of splenocytes and peripheric lymphocytes (CD4, CD8, B220 and CD45), and immunoglobulin responses by different white blood cells subpopulations (IgG, IgG1, IgG2a and IGM) (Supplementary figure 6).

**Figure 3.**
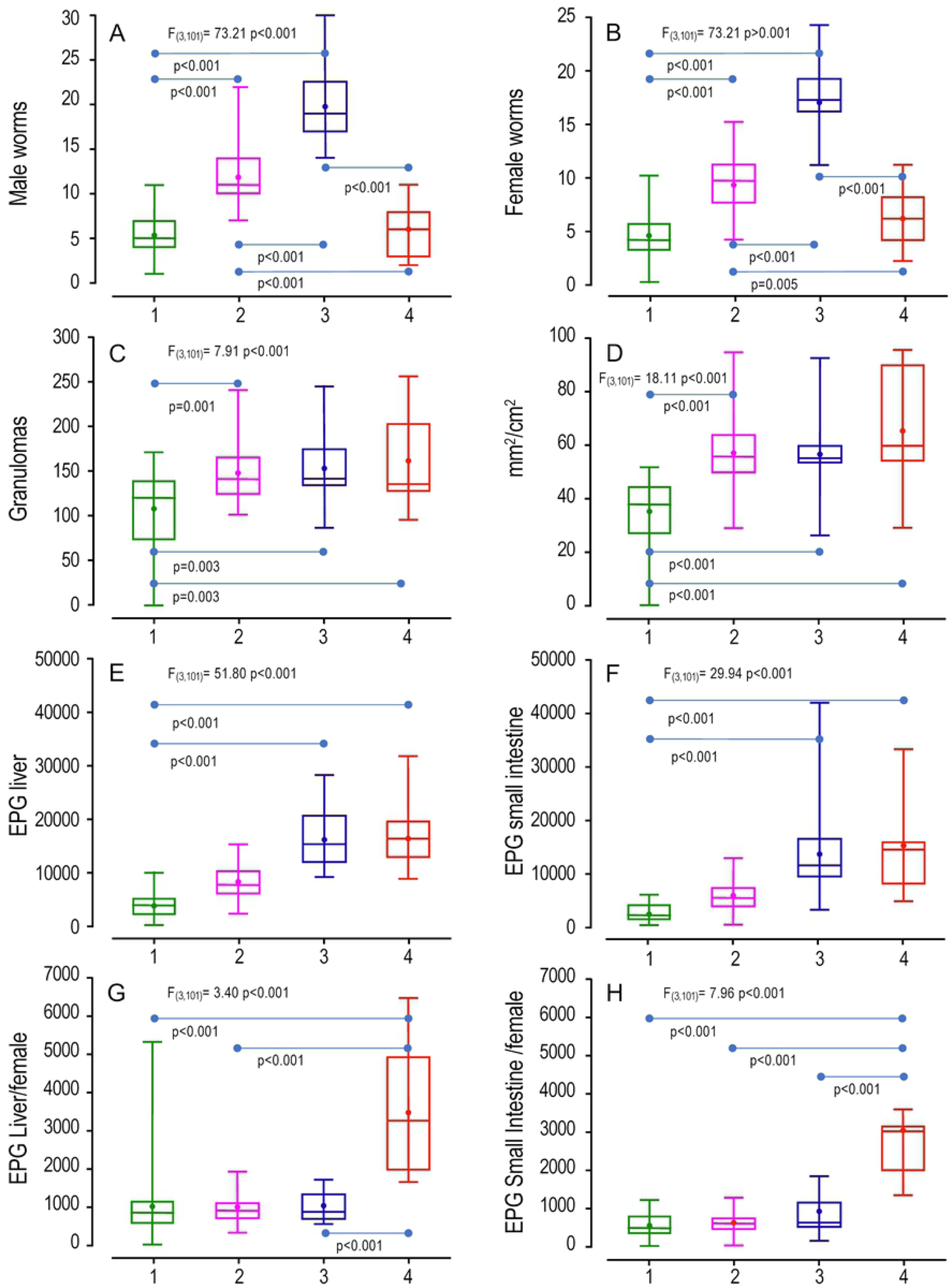
Differences in pathophenotypes and intermediate phenotypes between the four identified groups by k-means analysis. We evaluated the following variables: (A) male worms, (B) female worms (C) liver granulomas, (D) affected surface of the liver (mm^2^/cm^2^), (E) eggs per gram (EPG) of liver granulomas, (F) EPG of small intestine, (G) fecundity of females in liver (EPG Liver/female) and (H) fecundity of females in small intestine (EPG Small intestine/female). The comparison among the four groups was made by ANOVA. The comparisons between every two groups were made by the Tukey test.

Further, we studied medians in a heat chart where the maximum median of each variable within the four groups was taken as a reference (Figure 4). Data showed that Group 1 gathered mice with the lowest worm recovery, eggs in tissues and liver damage, thus, representing the less aggressive disease (Figure 3A-H) with the lowest medians (Figure 4). In Group 2 parasitological and pathological variables were significantly higher than Group 1 (Figure 3A-D), but eggs in tissues are similar to Group 1 (Figure 3E-H). Therefore Group 2 enclosed mice with a more aggressive disease. Group 3 showed a higher schistosome worm burden than Group 1 and 2 (Figure 3A and B) and significant liver damage, eggs in liver or eggs in intestine than Group 1 (Figure 3C-F), pointing a phenotype with more intense disease than Group 1 and 2, with the highest medians similar to Group 4 (Figure 4). Finally, Group 4 had the highest liver damage, counts in eggs in tissues and fecundity (Figure 3C-H, Figure 4) representing the most aggressive disease. Interestingly, however, the number of recovered worms was similar to Group 1 (Figure 3A and B).

**Figure 4.**
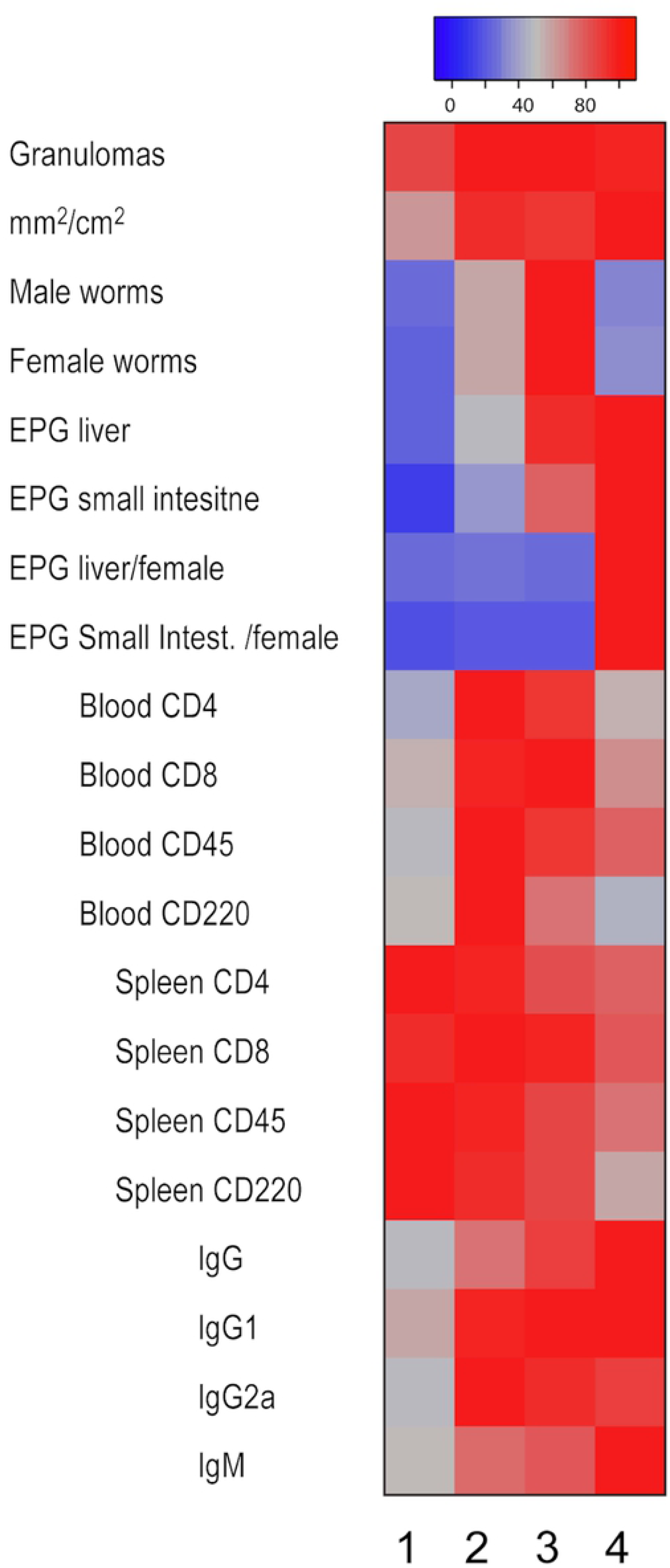
Correlation analysis of the medians of the variables between the four groups of mice. The maximum median of each variable within the four groups is taken as reference value. The proportion is represented, in percentage, that the median of each group has respect to the maximum, which indicates the variation experienced throughout the groups. The color represented when the difference between medium is low, it is red. Conversely, when the difference between medians is greater, the color represented is bluer.

For cell populations, no significant differences were observed between the four groups studied for the CD4 and CD8 populations, both in splenocytes or peripheral blood (supplementary Figure 6 A and B). Group 4 with the highest liver damage and eggs trapped in tissues, showed the lowest percentages in B220 and CD45 splenocytes (Supplementary Figure 6A). These differences were statistically significant in comparison with Group 1 and 2, and median analysis is concordance (Figure 4). In contrast, peripheral blood B220 and CD45 had no significant differences (Supplementary Figure 6B). Finally, we only found differences in total IgG antibodies, Group 3 and 4 showed more absorbance than Group 1 (Supplementary Figure 6C) with the highest medians (Figure 4). In the other hand, we did not find significant differences in IgG1, IgG2a or IgM levels between groups.

### Identifying chromosomal regions associated with susceptibility

The F1BX backcross cohort was genotyped with a platform of 961 informative SNPs and the quality control of the data was performed with *R/qtl* package R3.3.3. Genotypes of 92 mice were obtained, and the crossing was recognized as a backcross. After quality control, three SNPs and six mice with less than five or more than 20 overcrossings were eliminated, remaining a matrix with 86 individuals, 958 SNPs, and 20 phenotypes. We identied QTL genomic regions associated with the heterogeneous presentation of the disease (Figure 5) associated with parasite-pathological variables (Supplementary Table 1), lymphocyte subpopulations in peripheral bllod and spleen (Supplementary Table 2) and immunoglobulins (Supplementary Table 3). In our study, both the SNP rs6169611 on chromosome 15 and SNP rs13483259 of chromosome 18 were closely associated with the number of granulomas, which is the most representative variable of liver damage. The region including SNP rs6169611 in mouse chromosome 15 was found in human chromosome 8 (8q24.39 (Supplementary Figure 7A) and the region with SNP rs13483259 in mouse chromosome 18 was found in 18q11.2-12.3 h human chromosome 18 region.

**Figure 5.**
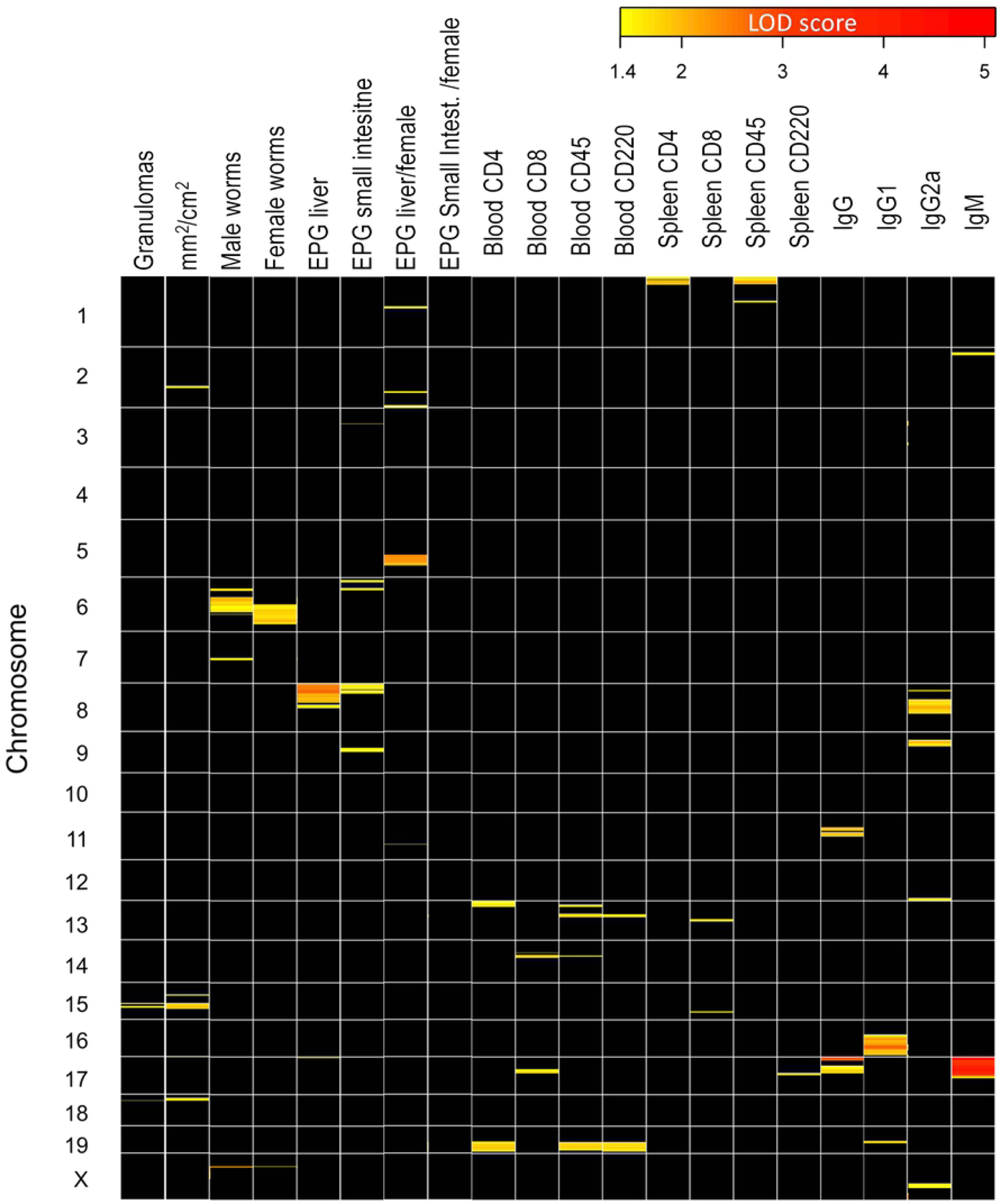
Heatmap of the chromosome QTL regions linked to pathophenotypes and intermediate phenotypes. The LOD score on an intensity scale from yellow to red represents the degree of linkage. QTLs with LOD scores lower than 1.4 were represented in black.

### Identification of genomic regions linked to disease susceptibility across the 4 groups identified in the F1BX

As the genetic component influences disease progression, we studied the QTLs associated with schistosomiasis by linkage analysis in the four defined groups in the F1BX cohort. We identified 11 loci associated with the disease out of 19 described above. Group 4 included a higher significant percentage of mice heterozygous for QTL11 (Chr 5), distal QTL5 (distal Chr 6) and QTL7 (Chr X). Regarding the eQTL3 (Chr2), the lowest percentage of heterozygotes was presented by Group 4, while in said QTL the highest percentage of heterozygosis was presented by Group 3. We also observed that for QTL8 (Chr 8) and QTL4 (Chr17), the groups with the highest percentage of heterozygous mice were 3 and 4, those with the most severe disease. Similarly, in the QTL4 (Chr17), Group 1 showed the lowest percentage of heterozygotes. Furthermore, in the distal eQTL5 in Chr6, Group 3 had the lowest number of heterozygous mice, whereas Group 4 showed the highest percentage of heterozygous mice (Figure 6).

**Figure 6.**
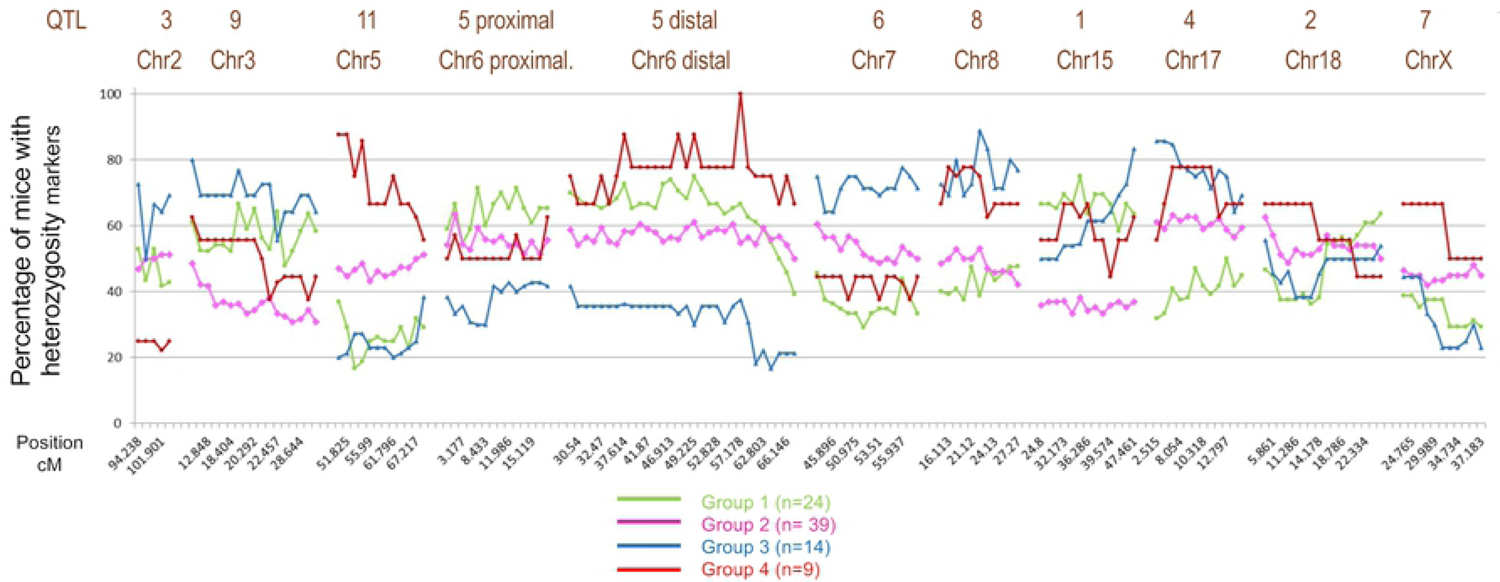
Comparison of the groups defined at the genetic level. The figure shows the percentage of mice with a particular heterozygous genetic marker, differentially present in the four groups of mice, according to the severity of the disease.

## Discussion

Only a minority of chronically exposed patients develops the most severe clinical form of schistosomiasis, severe hepatosplenic disease, periportal fibrosis and portal hypertension. Schistosomiasis is the second helminthiasis after soil-transmitted nematodes worldwide with high number of infected people [16] [17]. Hepatic granulomas and subsequent liver fibrosis vary significantly from individual to individual. Schistosome–host interaction is influenced by multiple factors, such as the type of immune response developed by the host, host genetic background, intensity, and number of infections. In endemic areas, high infection levels and severe periportal fibrosis are found concentrated in some families. Most of these differences do not reflect a dominant or recessive Mendelian inheritance of a single gene, but are under the control of multiple genes, defined as complex traits [18]. Susceptibility to infectious diseases responds to a model of complex traits, but the problem is that the same genotype can determine different phenotypes since complex traits have an incomplete penetrance, and they are subject to random effects and environmental [19]. There are publications where experimental *Schistosoma mansoni* infection is studied in different mouse strains, mostly made in 80s of the XX century[20–22]. These works analyze histopathological magnitudes, mainly granulomas and hepatic fibrosis, but do not identify susceptibility or infection resistance genetic markers.

In this study, we observed the schistosomiasis behavior in a genetically heterogeneous population obtained by backcrossing between a more susceptible (C57BL/6J) and a less susceptible (CBA/2J) mouse strains to generate a genetically and phenotypically heterogeneous F1BX cohort. Parasitological, pathological and magnitudes were studied to determine the level of infection, lesions and immune response [23]. The data obtained in F1BX mice showed worms recoveries like resistant parental strains. Also, eggs in tissues and liver damage were intermediate to the parental strains obtained in previous works in our research group [24]. When we compared F1BX mice phenotypic variable with F1B6CBA we observe a lower number of worms, both males and females, and a lower number of eggs per gram of liver [15]. After comparing F1BX males and females, we observed that there were no significant differences in the parasitological or pathological variables studied, indicating no association between the pathology caused by *S. mansoni* and the sex. Collectively, the data showed significant variation, with extreme values in the upper and lower limits, which is expected in the F1BX cohort. A correlation study between 20 phenotypic variables was performed. The parasitological and pathological magnitudes directly correlated between them, except for the number of worms and fecundity, whose relation is inverse. In this situation, with a more significant number of female worms, there is less production of eggs [25]. In the same way, immunological data were correlated between them, but there was a low correlation with parasitological or pathological magnitudes agreeing with an indirect impact of damages in the cell populations. The immune response against *S. mansoni* infection is very complex [23] since different types of cells of the immune system are involved, orchestrated by linker or effector molecules as immunoglobulins, which vary concerning the stage of the disease or the phase of the biological cycle [26].

We could define four groups or clusters of animals with different levels of disease according to the phenotypic variables analyzed employing Cluster Analysis dendrograms, Principal Component Analysis and k-means clusters. As expected in our model of phenotypic variability, eggs in tissues and hepatic damage had the most important loads determining the four groups ranging from mild to severe disease. This indicates that adult worms are not directly related to the histopathological lesions. Instead, eggs laying triggers the inflammatory reaction and granulomas formation. We observed that the cell subpopulations CD4 and CD8 were predominant in less and intermediate severity groups in both peripheral blood and spleen, decreasing drastically in the most severe group. CD4 lymphocytes are essential for granuloma formation and orchestrate the immune response against *S. mansoni* [27, 28] and CD8 lymphocytes modulate the Th2 immune response [29]. We observed the presence of antibodies production against the infection, but we could not observe any bias to Th1 or Th2 response associated with severity [30, 31]. The F1BX cohort showed wide variability from mild to severe infection intensity and disease. This variation makes it a suitable model for studing schistosomiasis in all ist gradation. Using this model, we could associate pathophenotypes and intermediate phenotypes to chromosomal regions with the help of SNPs.

Linkage analysis was performed using 958 differential SNPs with 86 mice and 20 variables after quality controls to detect genotyping errors, quantify crossovers and generate a suitable genetic map [32]. Once the recombination distances in the F1BX cohort were obtained, the QTL regions linked to the pathophenotypes were identified. We identified 19 chromosomal regions associated with the different phenotypic magnitudes studied with LOD-score above 1.4. The most exciting QTLs were QTL1 and QTL2 on chromosomes 15 and 18, respectively. The QTL1 was associated with chromosome 15 with three SNPs based on granulomas, affected liver surface, and spleen CD8 population. The QTL2 was associated with chromosome 18 by only one SNP with a high LOD score linked to granulomas and liver damage. The QTL3 and QTL4 were only associated with affected liver surface and IgM production and was located in chromosome 2 and 17. The QTL5 and QTL6 linked to chromosomes 6 and 8 were related to worm recovery and eggs in tissues and IgG2a.

Next, we looked for the syntenic regions of the QTLs identified by the Ensembl to study the homology between mice and humans. We chose for this study the QTL1 related to liver damage located on chromosome 15 with the marker peak SNP rs6169611 and eQTL2 located on chromosome 18 with the marker peak SNP rs13483259. We found syntenic regions with QTL1, within the human genome on chromosomes 5, 8, 12 and 22, particularly with a homologous region on human chromosome 8, in the 8q24.3 region. As QTL2, there were syntenic regions within the human genome on chromosomes 2, 5, 10 and 18, mainly on human chromosome 18, region 18q11.2-12.3. Previous studies in humans have shown an association with schistosomiasis susceptibility of specific chromosomal regions located on chromosomes 1, 5 and 6 [14]. Not all regions studied were associated with susceptibility to infection by *S. mansoni*, except SM2 locus (6q22-23) associated with liver fibrosis. Therefore, it can be deduced that the regions identified in this study in an experimental backcross model and with homology in the human genome have not been described in previous studies.

Finally, we analyzed the QTL association with the four clusters with different severity of schistosomiasis. We found that Group 4 (the most severe one) shows a high percentage of heterozygous mice for most SNPs that showed linkage. However, pathophenotypes are complex and manifest the effect of different intermediate phenotypes, which act at different levels of biological organization. For this reason, the results obtained did not collect all the genetic determinants that contributed to variability in the pathophenotypes. Linkage analysis of each phenotype revealed only those regions that contributed to the value of this with sufficient force. Genetic determinants of variability in intermediate phenotypes, showed no association with enough force to pathophenotypes could not be found by this analysis, and for this reason would be one of the causes of lost heritability [33]. Our study has limitations considering the number of mice included and the small size effect of involved chromosomal regions and the statistical power of QTL detection for the studied variables. Our results are the first steps towards identifying of genomic regions that control the phenotypic variation observed for *S. mansoni* infections. The dimension of significant regions will need additional fine mapping to propose reliable candidate genes influencing the defence response.

We can conclude that a F1BX cohort has been generated through backcross, with significant heterogeneity, helpful in studying the genetic influence on the susceptibility to infection produced by *Schistosoma mansoni*. The study has found 19 chromosomic regions spread in the mouse genome. Chromosomes 15 and 18 had the most relevant linkage to granulomas and hepatic surface damage, and it is homologous to regions in human chromosomes 8 and 18. These results will contribute to identifying target regions that control the variation of the complex parasite susceptibility trait in schistosomiasis and implementation these results in control and treatment schemes for especially susceptible populations.

## Materials and methods

### Animals and parasites

A backcross (F1BX) mouse cohort was generated after two stages; firstly, we crossed a mouse strain (CBA/2J) susceptible to schistosomiasis with a resistant one (C57BL/6J) to generate the F1B6CBA mice; secondly, the F1BX mice were generated by backcrossing. F1B6CBA female mice with CBA/2J males. Parental strains were purchased from Charles River Laboratories (Lyon, France) [34]. The animals’ procedures complied with the regulations on animal experimentation (L 32/2007, L 6/2013 RD 53/2013 and 2010/63/CE). The Ethics Committee of the University of Salamanca reviewed and approved the study (15/0018, 12/3351). All mice were housed in the Animal Research Facility of the University of Salamanca in standard conditions with free access to water and food *ad libitum*. The welfare and health of the animals were monitored during the experiment following the guidelines of the Federation of European Animal Science Laboratory Associations (FELASA). *S. mansoni* LB strain was routinely maintained on *Biomphalaria glabrata* snails and passages in CD1 mice every 2-3 months. Seven-weeks-old mice weighing 20-25 g were infected with cercariae of *S. mansoni*. The number and viability of the cercariae were determined using an Olympus SXZ9 stereomicroscope (Japan) [35].

### F1BX mouse cohort infection and parasitological and pathological traits obtained

F1BX mice were infected percutaneously with 150 ± 5 *S. mansoni* cercariae per mouse with the ring method and were euthanized nine weeks after the infection. Adult worms inside the portal and the mesenteric veins were recovered by perfusion with the help of an Olympus stereomicroscope SZX9 and a C-2000Z camera (Olympus Japan). Worms were obtained by opening the portal vein and retrieving them with tweezers, and the number of males and females, were recorded. The liver and intestine were removed, weighted and digested with 5% KOH overnight to determine the number of eggs in tissues. They were quantified in triplicate with a McMaster chamber. The worm’s fecundity was calculated by dividing the number of eggs in the liver and small intestine and the number of female worms [35]. Three photographs of different parts of each of mouse livers were collected before perfusion, to estimate the degree of tissue lesions. The number of granulomas per square centimeter was counted by two qualified researchers separately in each of the three micrographs. Adobe ImageJ software was used to enhance the contrast of images [36] and PowerPoint to set the sectors that facilitated the count. In case of dispute the intervention of a third investigator was requested.

### Peripheral and spleen white blood cell subpopulations quantified by flow cytometry

Blood samples were mixed with heparin, and splenocytes were obtained by aseptic prefusion of the spleen with 10 mL of sterile phosphate-buffered solution (PBS) at 37 °C. Samples of a minimum of 10^5^ peripheric white blood cells or splenocytes were mixed with 150 µl of lysis solution (154 mM NH_4_Cl, 10 mM KHCO_3_ and 0.082 mM EDTA (Sigma)) for 30 min, and then centrifuged at 237 *g.* The process was repeated, and the pellet was suspended in 150 μL of PBS with 2% fetal bovine serum (PBS-FBS). Monoclonal antibodies conjugated with fluorescein isothiocyanate (FITC) against CD4, with phycoerythrin (PE) against CD8, with allophycocyanin (APC) against B220 and with peridinin-chlorophyll proteins (PerCP-Cy™ 5.5) against CD45 from BD Pharmingen were diluted at 1/50 in PBS, 2% FBS, 25 μL per well and incubated for 30 min at room temperature in the dark. Then, cells were washed with 100 μl of PBS-FBS and cells were resuspended in 200 μL of PBS-FBS at 4° C in the dark. A flow cytometer FACSCalibur (BD Biosciences) was used and a minimum of 30,000 events were collected from each sample[37]. The data were analyzed by the free software Flowing Software 2.5.1 (http://flowingsoftware.btk.fi, Cell Imaging Core, Turku Centre for Biotechnology [38].

### Detection of anti-*S. mansoni* antibodies in mice infected with the parasite by ELISA

Serum samples were obtained before infection and necropsy to detect specific IgG, IgG1, IgG2a and IgM. Polystyrene Costar 96-well plates (Costar® 3596, Corning Inc) were upholstered with 5 μg/mL of SoSmAg (*S. mansoni* somatic antigen) in carbonate buffer pH 9.6 (100 μL/well) at 4 °C for 16 hr and then they were washed three times (200 μL/well) with PBS with 0.05% Tween 20 (PBST). Plates were blocked with bovine serum albumin (B4287, Sigma) 2% in PBST for 50 min at 37 °C (100 μL/well) and then they were washed three times with PBST. Sera were diluted 1:100 in PBST (100 μL/well), incubated for 1 h at 37 C in duplicate and washed three times with PBST as above. Conjugates with horseradish peroxidase anti-mouse IgGI, IgG1, IgG2a-HRP and IgM (Sigma) were used at 1:1000 in PBST (100 μL/well), incubated 1 hour at 37 °C and washed as above. Finally, they were developed with H_2_O_2_ (0.012%) and orthophenylenediamine (0.04%) in 0.1 M citrate buffer, pH 5.0 (100 μL/well). The reaction was stopped with 3N H_2_SO_4_ (50 μL/well) and read at 492 nm on a MultiSkan GO ELISA plate reader (Thermo Fisher Scientific, Vantaa, Finland) [35].

### SNP genotyping, quality control and linkage analysis

DNA was extracted from mouse tails using the DNeasy® Blood & Tissue Kit (Qiagen®, Hilden, Germany) and DNA concentration was measured with NanoDrop ND-1000 (Thermo-Fisher Scientific Inc.). Genotyping was carried out using the Illumina’s Mouse Low-Density Linkage Panel Assay at the National Genotyping Center (CeGen) of the National Center for Oncological Research (CNIO) [32]. A platform with 961 informative SNPs between C57BL/6J and CBA/2J strains was used. We prepared a matrix with 20 phenotypic variables with sex and genotype data. The autosome and female X homozygotes were coded as 0 and heterozygotes as 1. In males X chromosomes homozygous were coded as 0 and heterozygous as 2. Data quality control was carried out using the *R/qtl* package in R3.3.3 checking the crossing were recognized as a backcross, genotyping errors and quantifying crossovers. Also, the expected genetic map for F1BXmice was studied versus the estimated map. We performed a genetic linkage analysis using the pathophenotypes: parasitological and pathological variables, cellular populations and immunoglobulins. The genetic distance based on the recombination frequencies between markers in the F1BX cohort was compared to the genetic distance recorded for markers in the reference database Mouse Genome Informatics [8] http://cgd.jax.org/mousemapconverter. We used the method of maximum-likelihood mapping with the hope-maximization algorithm or HM algorithm. The Haldane function was used to calculate the conditional genotype, with a step size of 2.5 cM and a genotyping error of 0.001 [39]. We used the LOD-score to calculate the statistical significance of the linkage of the QTLs found. We use the criteria established by Lander and Kruglyak [40] to consider the significance level of association observed between a QTL and the phenotype of interest. Therefore, values of LOD-score higher than 1.4 were considered to indicate the presence of suggestive linkage. The Ensembl bioinformatics tool (https://www.ensembl.org/index.html) was used to identify syntenic regions between mouse and human.

## Data analysis

Mean and standard error of the mean were determined in each variable and normal distribution was checked with the Kolmogorov-Smirnov test. ANOVA followed by *post-hoc* Tukey’ test or Student t-test were employed to identify differences between groups. The Pearson correlation coefficient (r) was used to study the dependence of two variables and the statistical significance was performed using the Student t-test.

Multivariant models were generated to study mice’s similarities and consider sex influence in worm recovery, eggs in the liver and intestine, fecundity, granulomas and the affected liver surface. All the variables were standardized to 0 mean and standard deviation to 1 to avoid the magnitude differences of the variables. Cluster analysis (CA) dendrogram was used to evidence the distances between the cases. Principal components analysis (PCA) extracts condensed information in a few variables, identifing affinity groups and the influence of each variable. With the information obtained with CA [41] and PCA [42] we applied the algorithm of conglomerates by k-means to generate groups or clusters according to the degree of severity observed in mice. Further analyses were performed in the groups obtained including median proportions in each variable. A significant difference was considered in all tests when the p-value for accepting the null hypothesis was less than 0.05. The results were analyzed using the SIMFIT statistical package Windows version 7.3.7 [43].

## Fundings

This research was funded by RICET RD16/0027/0018.

## Contributors

Conceptualization: J.H-G., J.L-A., J.P-L. and A.M.; data curation: J.H-G., J.L-A., A.B-G., F-J.B.; formal analysis: J.H-G., J.L-A., A.B-G., F-J.B. and J.P-L.; funding acquisition: A.M.; investigation: J.H-G., J.L-A., A.B-G. and B.V.; methodology: J.H-G., J.L-A, A.B-G., B.V. and F-J.B.; project administration: J.P-L. and A.M.; resources: J.L-A., B.V., J.P-L. and A.M.; software: J.H-G., J.L-A., A.B-G., F-J.B.; supervision: J.H-G., J.L-A., A.B-G., B.V. and F-J.B.; validation: J.H-G. and J.L-A.; visualization: J.H-G. and J.L-A.; writing – original draft: J.H-G. and J.L-A.; writing – review and editing: J.P-L. and A.M.

---

**Supplementary figure 1.**
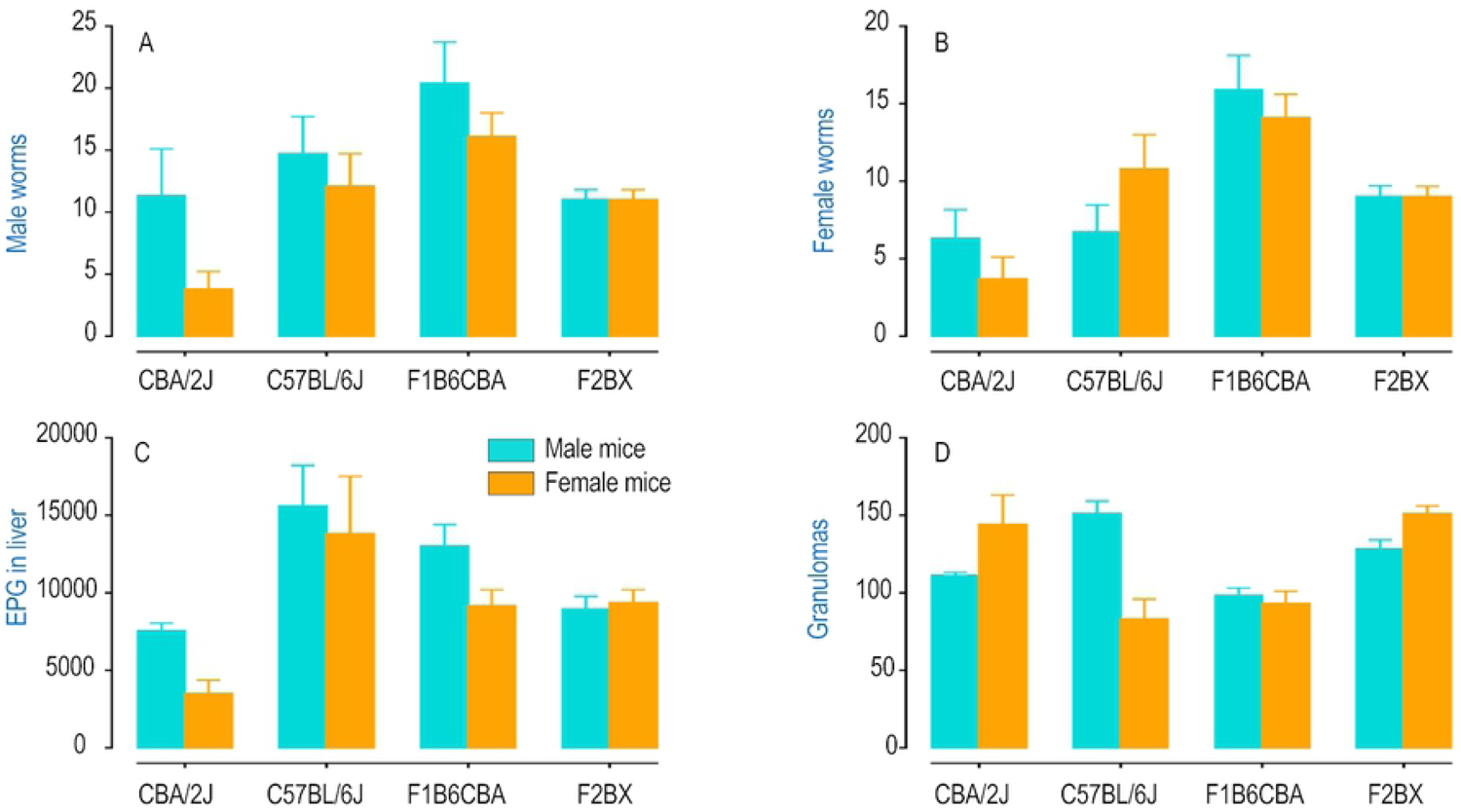
Representation of parasitological variables (A number of recovered male worms and B number of recovered female worms) and pathological (C eggs per gram of liver and D number of granulomas /cm2 of liver surface) between parental strains (CBA/2J, C57/2J) F1 progeny (F1B6BCA) and Backcross (F1BCX). The means and the standard error of the mean (SEM) were represented.

**Supplementary figure 2.**
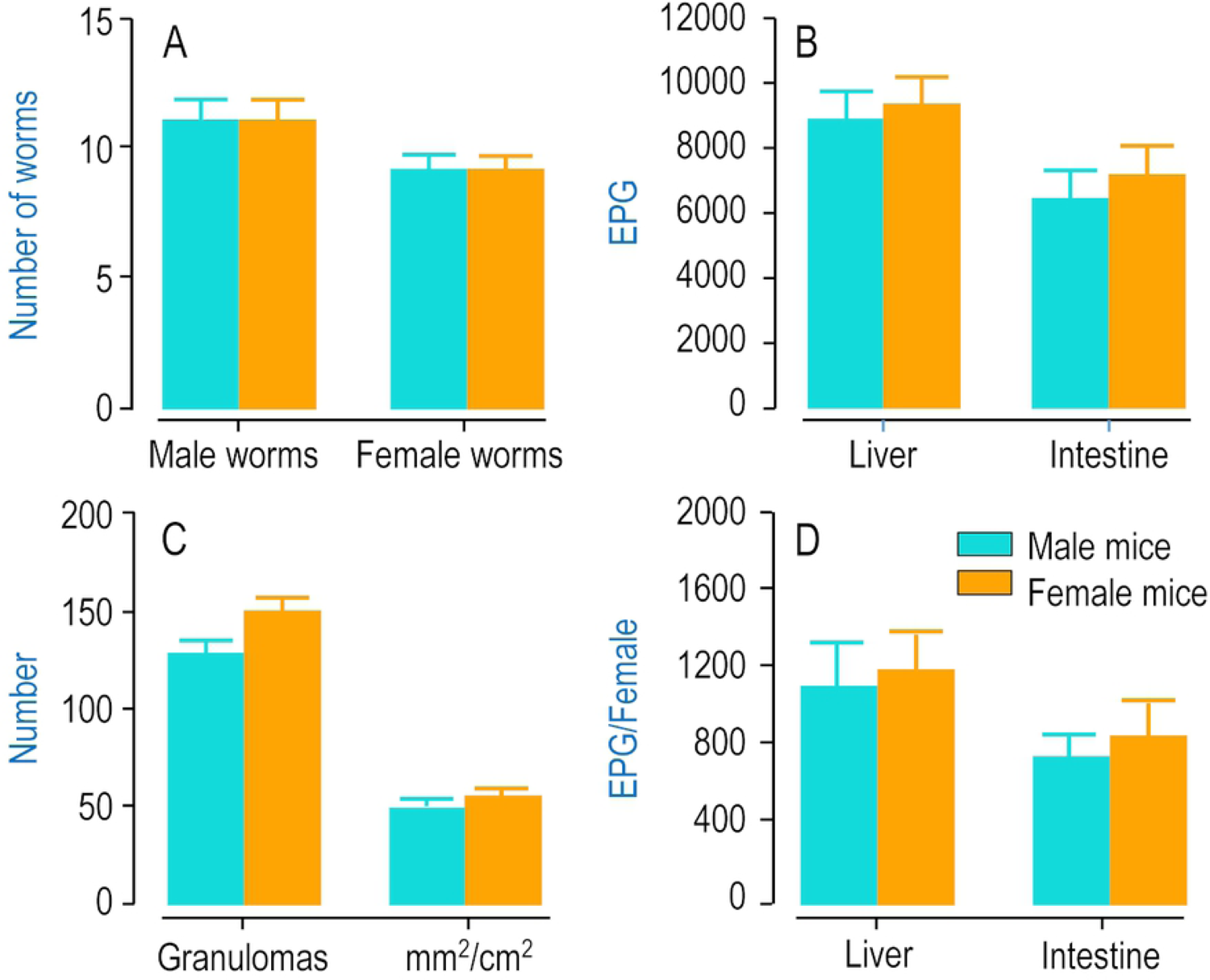
Comparison betweenF1BX male and female mice: (A) number of recovered male and female worms, (B) number of eggs per gram (EPG) of liver and small intestine, (C) number of granulomas and affected surface of the liver and (D) fecundity of female worms in liver and small intestine. The means and the standard error of the mean (SEM) were represented.

**Supplementary figure 3.**
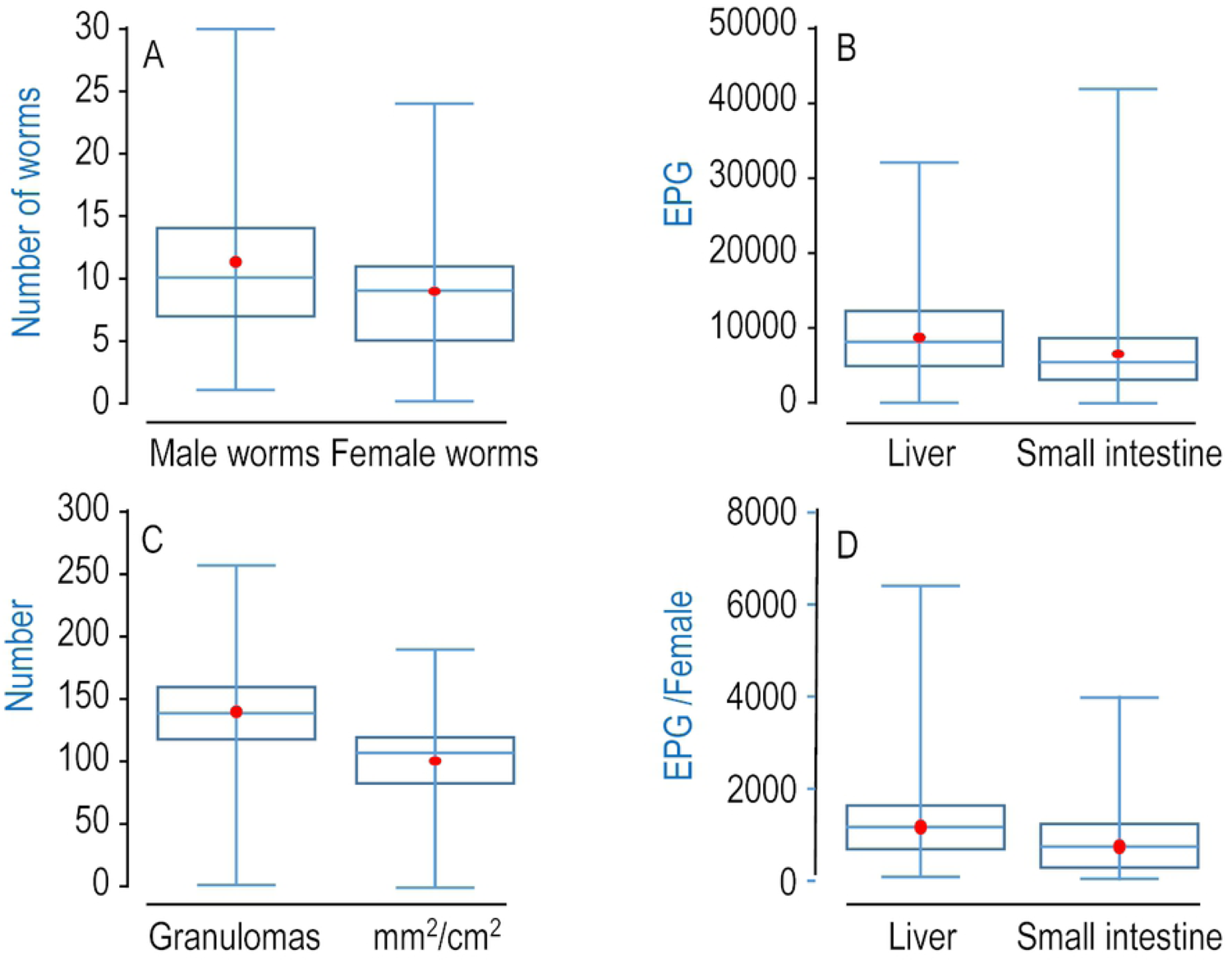
An exploratory study of parasitological variables: A number of recovered male and female worms; B Number of eggs per gram (EPG) of liver and small intestine; C number of granulomas and affected liver surface (mm/cm2) of) for the F1BX cohort. Box plot and the mean as a red circle.

**Supplementary figure 4.**
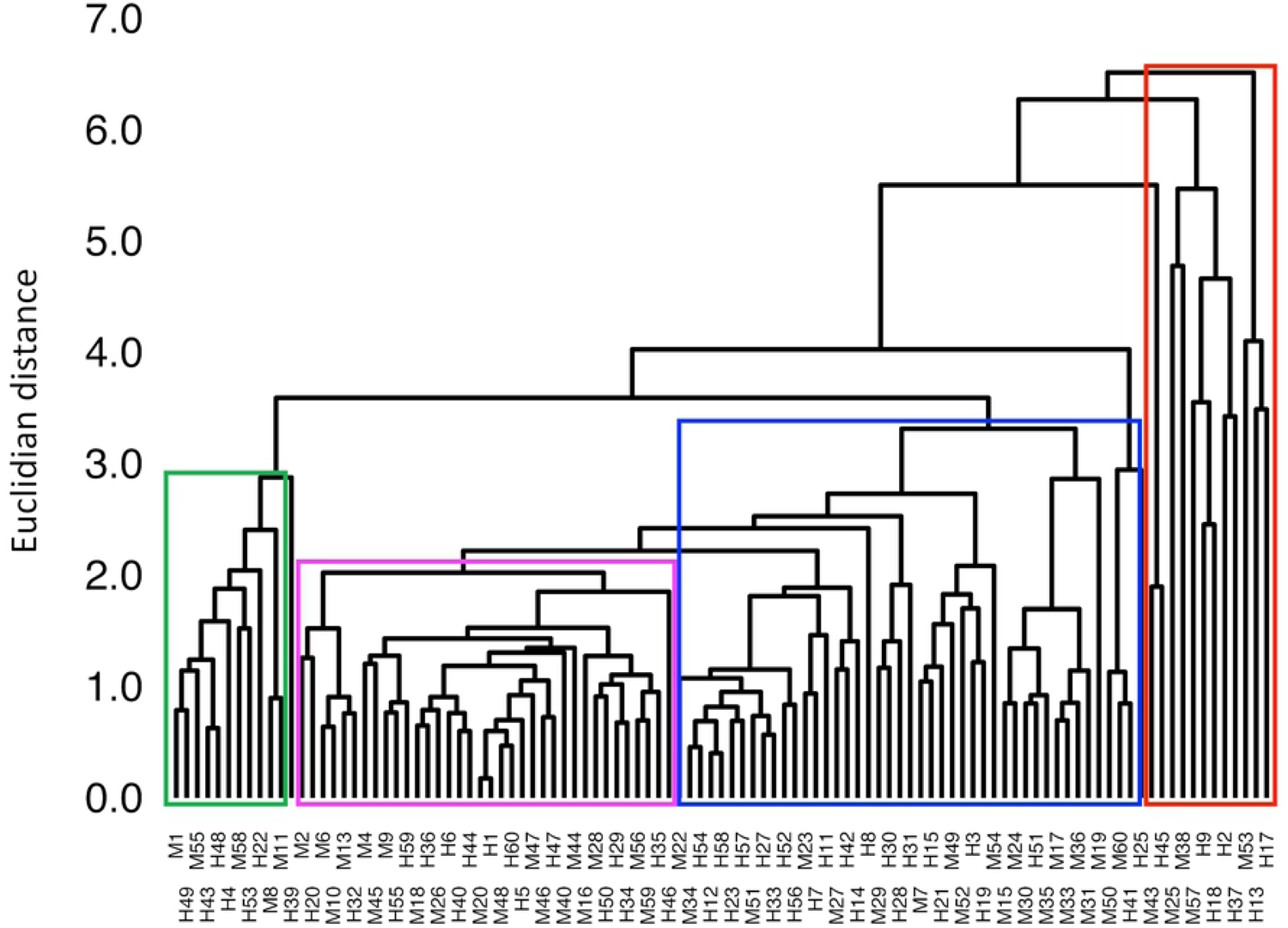
Dendrogram with all parasitological and pathological variables.

**Supplementary figure 5.**
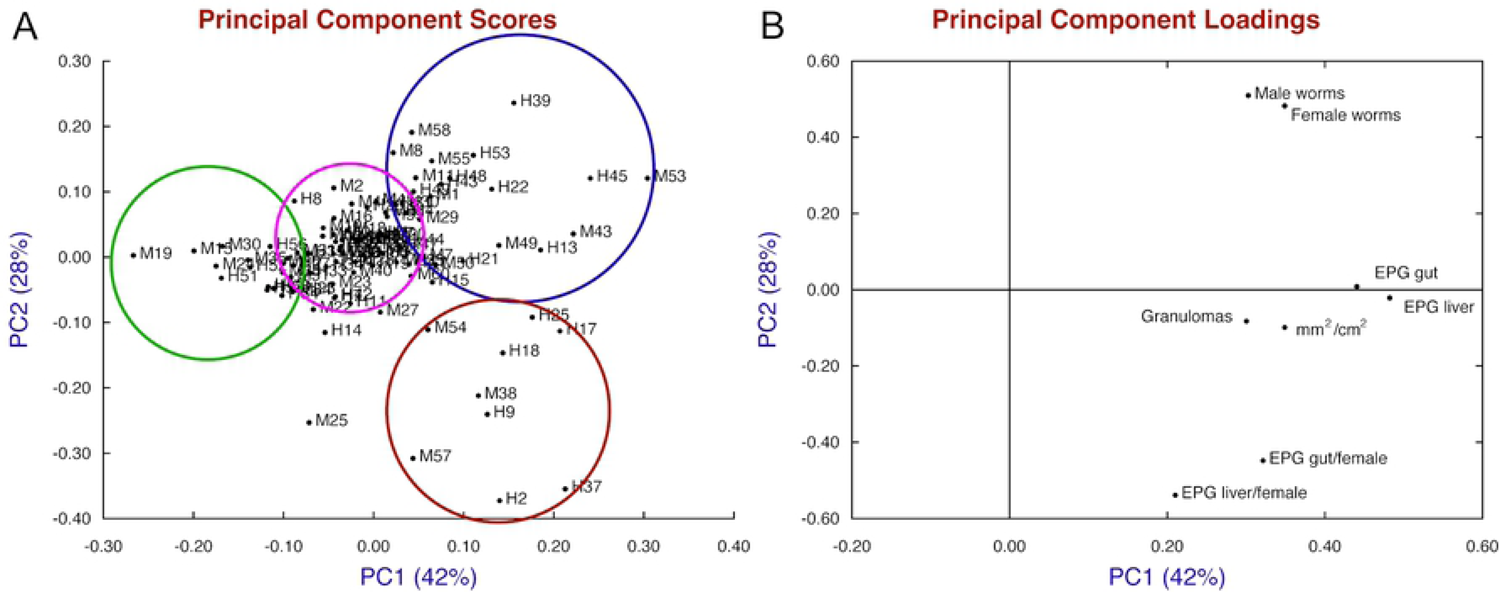
Principal components analysis (PCA). A) Representation of the scores with four tentative groups of mice in the F1BX cohort. B) Representation of variable loadings.

**Supplementary figure 6.**
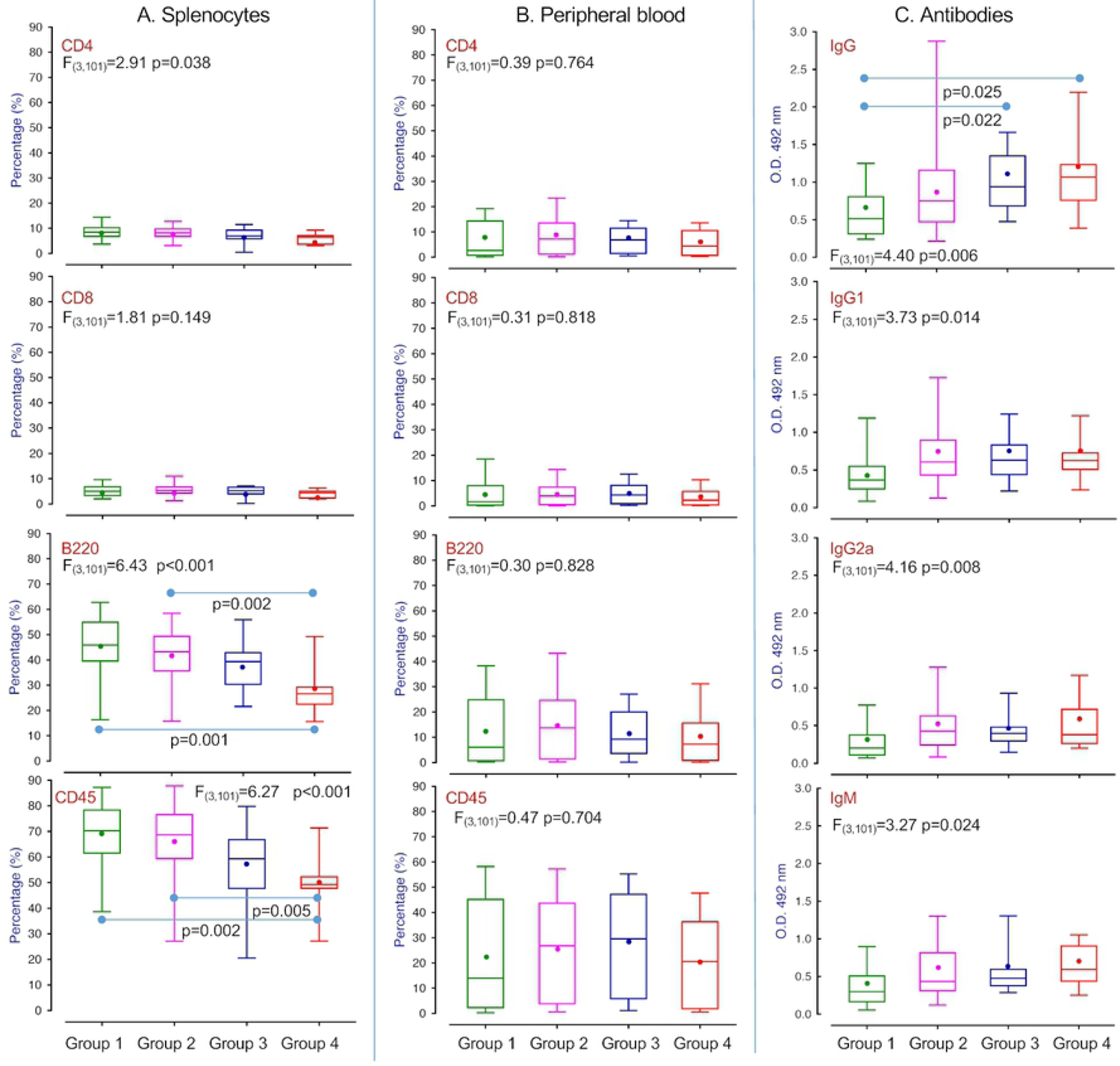
Differences between the four groups of schistosomiasis degree regarding cell lymphocyte subpopulations in peripheral blood: (A) CD4, (B) CD8, (C) B220 and (D) CD45, cell subpopulations in splenocytes: (E) CD4, (F) CD8, (G) B220 and (H) CD45 and immunoglobulins (I) IgG, (J) IgG1, (K) IgG2a and (L) IgM nine-week postinfection. The comparison among the four groups was made by ANOVA, and the Tukey test was made between every two groups.

**Supplementary Figure 7.**
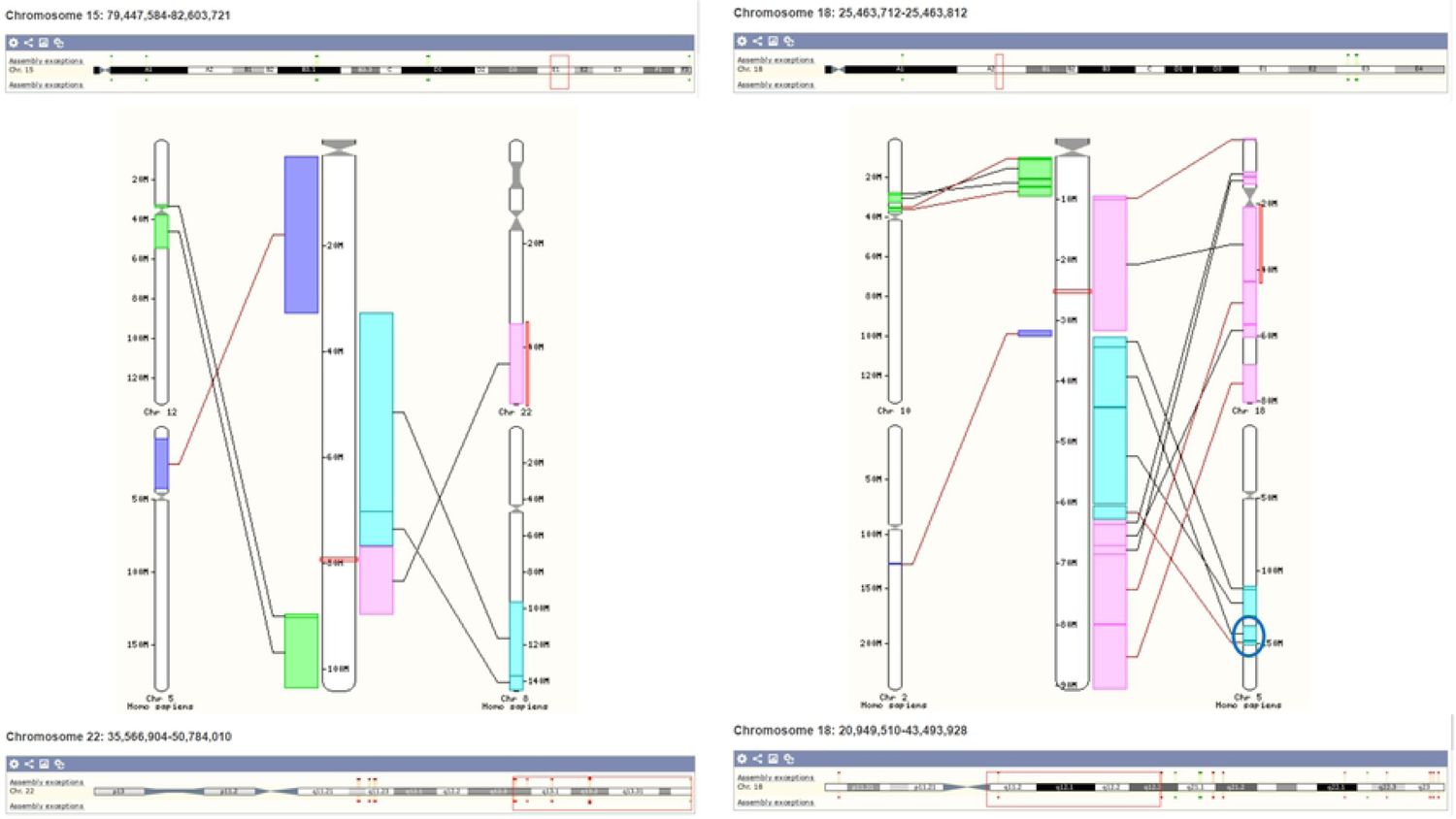
Syntenic regions located in humans from identified quantitative trace loci (QTL) in mice. A) The QTL1 on mouse chromosome 15 was mainly located on human chromosome 8. B) The QTL2 on mouse chromosome 18 was mainly located on human chromosome 18.

**Supplementary Table 1.** Quantitative trace loci (QTL), associated chromosome (Chr) and location (cM) with LDO scores, and the average of the variables in homozygous (AA) and heterozygous (AB) genotypes and the allele combination with higher effect in parasitological and pathological phenotypes in the backcross cohort F1BX. (EPG-Eggs per gram).

| Phenotype | eQTL | Chr. | Marker Peak | Location (cM) | LOD Score | Average ± SEM | | | | Higher Effect Allele |
| --- | --- | --- | --- | --- | --- | --- | --- | --- | --- | --- |
|  |  |  |  |  |  | AA | AB | AY | BY |  |
| Granulomas | eQTL1 | 15 | rs6169611 | 32.17 | 1.51 | 152.61±6.49 | 127.64±6.64 | | | AA |
| | eQTL2 | 18 | rs13483259 | 14.18 | 1.48 | 129.29±6.40 | 153.78±6.70 | | | AB |
| Injured hepatic surface | eQTL3 | 2 | rs13476772 | 63.25 | 1.47 | 56.63±2.76 | 46.70±2.52 | | | AA |
| | eQTL1 | 15 | rs13482664 | 37.72 | 2.03 | 56.84±2.56 | 45.31±2.62 | | | AA |
| | eQTL4 | 17 | rs13482845 | 2.52 | 1.54 | 45.59±2.80 | 55.65±2.49 | | | AB |
| | eQTL2 | 18 | rs13483259 | 14.18 | 1.54 | 46.59±2.55 | 56.78±2.80 | | | AB |
| Male worms | eQTL5 | 6 | rs13478799 | 30.04 | 2.01 | 13.11±0.92 | 9.31±0.80 | | | AA |
| | eQTL6 | 7 | rs13479382 | 51.74 | 1.64 | 9.31±0.84 | 12.73±0.88 | | | AB |
| | eQTL7 | X | rs13483769 | 30.62 | 2.28 | 12.81±1.20 | 10.46±1.08 | 13.07±1.29 | 7.74±1.22 | AY/AA/AB/BY |
| Female worms | eQTL5 | 6 | c6.loc57.5 | 59.38 | 2.11 | 10.51±0.73 | 7.28±0.70 | | | AA |
| | eQTL7 | X | rs13483769 | 30.62 | 1.45 | 9.67±1.03 | 8.15±0.93 | 10.73±1.10 | 7.07±1.05 | AY/AA/AB/BY |
| EPG liver | eQTL8 | 8 | rs13479680 | 19.01 | 2.82 | 6974.25±885.31 | 11560.76±844.22 | | | AB |
| | eQTL4 | 17 | rs13474581 | 4.55 | 1.41 | 7481.65±966.22 | 10806.63±839.62 | | | AB |
| EPG gut | eQTL9 | 3 | rs13477093 | 20.78 | 1.67 | 5162.24±809.89 | 8584.90±892.88 | | | AB |
| | eQTL5 | 6 | rs3023064 | 7.99 | 1.50 | 8424.47±893.53 | 5169.28±849.21 | | | AA |
| | eQTL8 | 8 | rs13479680 | 19.01 | 1.61 | 4967.71±869.99 | 8301.83±831.33 | | | AB |
| EPG of liver/female | eQTL10 | 1 | rs13475979 | 47.01 | 1.49 | 1685.07±199.20 | 1031.32±142.09 | | | AA |
| | eQTL3 | 2 | rs6257221 | 102.73 | 1.56 | 1552.30±159.78 | 922.42±167.33 | | | AA |
| | eQTL11 | 5 | rs13478485 | 60.10 | 2.37 | 954.78±142.56 | 1752.94±185.19 | | | AB |
| | eQTL12 | 11 | rs13481156 | 55.29 | 1.41 | 967.27±159.11 | 1573.58±169.49 | | | AB |
| EPG of intestine/female | -- | - | - | - | - |  |  |  |  |  |

**Supplementary Table 2.** Quantitative trace loci (QTL), associated chromosome (Chr), location (cM) with LOD scores and the average of the variables in homozygous (AA) and heterozygous (AB) genotypes and the allele combination with higher effect in cell populations at nine-week (W9) post-infection in the backcross cohort F1BX.

| Phenotype | eQTL | Chr. | Marker Peak | Location (cM) | LOD Score | Average ± SEM | | | | Higher Effect Allele |
| --- | --- | --- | --- | --- | --- | --- | --- | --- | --- | --- |
|  |  |  |  |  |  | AA | AB | AY | BY |  |
| CD4 Blood W9 | eQTL13 | 13 | rs13481740 | 13.23 | 1.68 | 5.77±0.83 | 9.13±0.84 | | | AB |
| | eQTL14 | 19 | rs13483665 | 45.23 | 1.95 | 9.34±0.86 | 5.73±0.80 | | | AA |
| CD8 Blood W9 | eQTL15 | 14 | rs8251327 | 28.08 | 1.81 | 7.56±0.73 | 4.42±0.78 | | | AA |
| | eQTL4 | 17 | rs13482998 | 20.63 | 1.92 | 7.92±0.80 | 4.67±0.71 | | | AA |
| CD45 Blood W9 | eQTL13 | 13 | rs13481826 | 30.69 | 1.86 | 30.27±4.21 | 49.03±4.62 | | | AB |
| | eQTL15 | 14 | rs8251327 | 28.08 | 1.44 | 46.47±4.61 | 29.93±4.57 | | | AA |
| | eQTL14 | 19 | c19.loc40 | 44.08 | 1.85 | 48.68±4.57 | 30.17±4.26 | | | AA |
| B220_Blood_W9 | eQTL13 | 13 | rs13481826 | 30.69 | 1.69 | 11.65±1.80 | 19.30±1.98 | | | AB |
| | eQTL16 | 19 | rs13483665 | 45.23 | 1.83 | 19.42±1.96 | 11.41±1.81 | | | AA |
| CD4 Spleen W9 | eQTL10 | 1 | rs13475740 | 4.95 | 2.03 | 6.92±0.39 | 8.56±0.35 | | | AB |
| CD8 Spleen W9 | eQTL13 | 13 | rs4229866 | 35.76 | 1.68 | 4.69±0.29 | 5.97±0.33 | | | AB |
| | eQTL1 | 15 | rs13482696 | 43.56 | 1.49 | 5.81±0.30 | 4.65±0.31 | | | AA |
| CD45 Spleen W9 | eQTL10 | 1 | rs13475749 | 6.03 | 2.10 | 58.42±2.34 | 68.29±1.99 | | | AB |
| B220 Spleen W9 | eQTL4 | 17 | rs13483013 | 24.72 | 1.64 | 42.45±2.56 | 32.61±2.30 | | | AA |

**Supplementary Table 3.** Quantitative trace loci (QTL), associated chromosome (Chr), location (cM) with LOD scores and average of the variables in homozygous (AA) and heterozygous (AB) genotypes and the allele combination with higher effect in circulant antibodies at nine-weeks (W9) post-infection in the backcross cohort F1BX

| Phenotype | eQTL | Chr. | Marker Peak | Location (cM) | LOD Score | Average ± SEM | | | | Higher Effect Allele |
| --- | --- | --- | --- | --- | --- | --- | --- | --- | --- | --- |
|  |  |  |  |  |  | AA | AB | AY | BY |  |
| IgG W9 | eQTL12 | 11 | rs13481062 | 38.91 | 2.03 | 0.7±0.07 | 1.02±0.07 | | | AB |
| | eQTL4 | 17 | rs13482845 | 2.52 | 3.42 | 0.15±0.01 | 0.17±0.009 | | | AB |
| IgG1 W9 | eQTL17 | 16 | rs4204623 | 42.53 | 2.85 | 0.48±0.04 | 0.70±0.04 | | | AB |
| | eQTL16 | 19 | rs13483612 | 32.11 | 1.74 | 0.49±0.04 | 0.67±0.04 | | | AB |
| IgG2a W9 | eQTL8 | 8 | rs13479846 | 37.83 | 2.13 | 0.31±0.036 | 0.48±0.037 | | | AB |
| | eQTL18 | 9 | rs4227609 | 24.58 | 2.31 | 0.31±0.035 | 0.48±0.036 | | | AB |
| | eQTL19 | 12 | c12.loc57.5 | 59.92 | 1.54 | 0.46±0.037 | 0.32±0.037 | | | AA |
| | eQTL7 | X | rs13483959 | 49.16 | 1.44 | 0.33±0.047 | 0.52±0.052 | 0.34±0.055 | 0.37±0.053 | AB/BY/AY/AA |
| IgGM W9 | eQTL3 | 2 | c2.loc20 | 23.68 | 1.57 | 0.61±0.045 | 0.44±0.044 | | | AA |
| | eQTL4 | 17 | rs13482845 | 2.52 | 4.96 | 0.37±0.030 | 0.67±0.047 | | | AB |

## Notes

### Competing Interest Statement

The authors have declared no competing interest.

